# Sexual dimorphism in zymosan-induced arthritis is linked with a higher IFN response in myeloid cell subsets that faithfully recapitulate RA synovial cell clusters

**DOI:** 10.1101/2025.11.24.690174

**Authors:** Richard D Bell, Mary Huang, Ruoxi Yuan, Claire Wingert, Bikash Mishra, Seda Seren, Toolika Singh, Susan MacLauchlan, Ellen Gravellese, Anvita Singh, Amit Lakhanpal, Anne Bass, Laura B. Donlin, Lionel B Ivashkiv

## Abstract

A key feature of rheumatoid arthritis (RA) is sexual dimorphism, with a higher incidence of RA in females. Myeloid cells are key drivers of the inflammatory effector phase of RA, but little is known about their contribution to a sexually dimorphic arthritis phenotype. We wished to utilize an arthritis model that recapitulates RA pathogenic myeloid cell subsets defined by single cell transcriptome analysis to investigate sex differences in human disease-relevant cell types and related molecular pathways. We developed a computational strategy that rigorously maps scRNAseq-defined mouse arthritis myeloid cells and clusters onto previously defined RA subsets. Synovial myeloid cells in the zymosan-induced arthritis (ZIA) model closely recapitulated four pathogenic RA myeloid cell subsets, strongly supporting the disease-relevance of the ZIA model. ZIA also effectively modeled myeloid cells in immune checkpoint inhibitor arthritis (ICI-A). These RA, ICI-A and ZIA myeloid cells express genes in TNF-NF-kB, PGE2, and IFN-STAT pathways. In ZIA, an induction phase dominated by cell clusters expressing NF-kB and PGE2 pathways transitioned to peak arthritis characterized by cell subsets co-expressing NF-kB and IFN pathways, as occurs in RA and various inflammatory diseases. Female mice exhibited increased arthritis linked with early induction of interferon-stimulated genes and increased expansion of IFN signature-expressing cell subsets. Our study identifies a computational strategy and animal model that enables investigation of recently described pathogenic myeloid cell subsets, and links increased arthritis in female mice with specific myeloid cell subsets that exhibit a stronger IFN response.

## Introduction

Rheumatoid arthritis (RA) is an autoimmune disease characterized by inflammation of synovial joints (synovitis) that leads to swelling, pain and destruction of cartilage and bone^1^. Autoimmunity and production of autoantibodies against citrullinated peptides and IgM precede the development of overt synovitis. Synovitis is characterized by infiltration of joints by immune cells including T cells, B cells/plasma cells and myeloid cells (monocytes, macrophages, DCs and synovial fluid neutrophils), and proliferation of synovial fibroblasts that exhibit an inflammatory and tissue-destructive phenotype^1^. Synovial monocytes and macrophages (MonoMacs) are major producers of pathogenic cytokines such as TNF and IL-1b, and have been closely linked with synovitis pathogenesis^2^. Recent work has demonstrated that RA synovial CD4+ and CD8+ T cells highly express IFN-g (but not IL-4 or IL-17)^3–5^, and corroborated decades of work showing pervasive expression of interferon-stimulated genes (‘IFN signature’) in multiple synovial cell types including myeloid cells, B cells, and fibroblasts. Similar to RA, arthritis that occurs after immune checkpoint inhibition (ICI-A) is characterized by immune infiltrates, activated monocytes and IFN-g-expressing T cells^6^. This work highlights the importance of myeloid cells and IFN-g in the effector inflammatory phase of arthritis, including RA and ICI-A.

Synovial myeloid cells play a critical role in joint homeostasis^2,7^, modulating persistent inflammation and joint destruction^8^, and mediating response to therapy^3,9^ in RA. The recent explosion in knowledge about RA synovitis at the single cell and molecular levels provides a great opportunity to test which cell subtypes and molecular pathways are causal and pathogenic^4,10–14^. Putative pathogenic cell populations include macrophages that express inflammatory factors and EGFR ligands which stimulate fibroblasts to be invasive and tissue destructive (IL1B+ HBEGF+ Macs^3,4,8,14,15^) and monocyte-macrophages (MoMacs) that express interferon-stimulated genes (ISGs) that encode lymphocyte recruiting and stimulatory chemokines/cytokines (STAT1+CXCL10+^4,15^). Although determining the importance of these cells and pathways in the pathogenesis of RA will ultimately require clinical trials, one important next step is to utilize the new multi-omic data to inform the use of animal models to more closely mimic key aspects of human RA and test causality of proposed pathogenic mechanisms to inform future clinical trials. Animal models that recapitulate pathogenic RA myeloid subsets will enable testing of their roles in distinct pathogenic processes such as tissue destruction and immune cell recruitment, and may lead to a better understanding of the pathogenesis of RA.

Commonly used pre-clinical models of RA can be broadly divided into models of induced autoimmunity against joint components such as collagen (e.g. antibody-dependent), and models of the effector phase of joint inflammation (synovitis), typically induced by injection of antibodies against joint components or transgenic expression of TNF (e.g. antibody-independent)^16–18^.

Although these models capture the tissue destructive arthritic phenotype and have been useful for preclinical testing of therapeutic agents, there are several limitations in their ability to mimic RA. The induced autoimmunity models are typically driven by Th17 cells, whereas RA has minimal synovial Th17 cells or IL-17, is not responsive to IL-17 blockade, and is generally considered not to be a Th17 disease^17,19–23^. It is difficult to dissect mechanisms that regulate effector phase synovitis in autoimmune models, as perturbations of cell and molecular pathways affects both the development of autoimmunity and synovitis. Many models of the inflammatory effector phase do not capture the lymphocyte-driven component of synovitis, and to our knowledge none of these models recapitulate the T cell-IFN-g contribution to synovitis.

Additionally, there is minimal information about how well these models recapitulate the proposed pathogenic myeloid subsets that have been described in RA. Utilization of a model of synovitis that incorporates an IFN response and recapitulates key synovial myeloid subsets would increase our ability to obtain mechanistic insights into arthritis pathogenesis.

An additional, but rarely modeled, key feature of rheumatoid arthritis (RA) is its pronounced sexual dimorphism: women are affected approximately three times more often than men (2:1 ratio)^24–26^. This increased prevalence contributes substantially to the healthcare burden, as women with RA have the second highest incidence among all autoimmune diseases^26^. Moreover, women tend to exhibit higher disease activity^27,28^ and achieve remission less frequently than men^28^, accounting for up to 92% of treatment-refractory patients^29^. Despite recognition of these sex differences in RA for more than 30 years^30^, mechanistic understanding remains limited, and no sex-specific therapies have yet been developed to either harness the apparent protection observed in males or mitigate the heightened susceptibility in females.

While our understanding of sex-specific immune regulation has made great strides in the last 10 years, specifically in understanding the TLR7-X-chromosome inactivation axis^31^, the role of androgens limiting ILC2-dependent dendritic cell activation in skin immunity^32^, the T-regulatory 2cell response to sex hormones^33^, and the myeloid cell transcriptional differences in IFN response^34,35^, there is a lack of understanding of how these processes specifically contribute to RA pathogenesis^36^. This is largely hampered by the lack of murine models that recapitulate the features of sexual dimorphism seen in humans. The few mechanistic studies suggest roles for B-cells^37–39^, T-cells^40–43^ and osteoclasts^44^ in contributing to the female specific disease phenotype, while a sexually dimorphic role for myeloid effector cells in the inflammatory component of disease is as yet undescribed.

We wished to investigate sexual dimorphism in the role of myeloid cells in synovitis. We reasoned that an optimal model would focus solely on the effector phase of arthritis to avoid confounding effects of sexual dimorphism in adaptive immunity, and would recapitulate key aspects of RA including immune cell infiltrates, an IFN signature, and most importantly key pathogenic myeloid subsets identified by single cell profiling. We chose zymosan induced arthritis (ZIA) which is an acute arthritis model that is independent of prior generation of autoimmunity, histologically recapitulates the immune cell infiltrates and tissue destructive nature of RA^45^, exhibits strong activation of STAT1 (indicative of IFN activity)^16–18,46^ and, similar to RA, is partially responsive to TNF and IL-6 blockade therapy^46,47^. We developed a computational strategy to rigorously map ZIA myeloid cells and cell clusters onto RA and ICI-A pathogenic cell subsets. We show that ZIA myeloid cells closely recapitulate pathogenic RA and ICI-A myeloid cell subsets, strongly supporting the relevance of using ZIA as an arthritis model. We found that ZIA is sexually dimorphic and linked increased arthritis in female mice with an earlier and stronger IFN response in specific myeloid cell subsets.

## Results

### Zymosan induces sexually dimorphic synovial arthritis

We first performed a dose response of intra-articularly injected zymosan that demonstrated robust induction of ZIA by 180 mg of zymosan, which established zymosan doses that we used in our study (**Supplemental Figure 1A-C**). We then investigated the role of sex on arthritis development by performing a longitudinal study of ZIA in C57BL/6 mice. Injected knees of female mice were consistently more swollen than ZIA knees of male mice over the 4-6 week course of active arthritis (**Figure 1A**). Measurement of the area under the arthritis curve (AUC) showed that differences between females and males were highly significant (p = 0.0076, **Figure 1A**). These female-male differences were corroborated by histology using an unbiased computer vision and machine learning approach (CPath-Arthritis)^48^ to measure synovial area and cell counts (**Figure 1B, C**). Synovial area and cell counts showed increased arthritis in female mice that was most pronounced on D14 and persisted through D28. Female mouse synovial area and cell counts increased from D7 to D14, whereas male mouse synovial area and counts did not significantly change between D7 and D14. Differences between arthritis in female and male mice are further illustrated by representative histology images shown in **Figure 1D**. Humanized TLR8 (Tg8) transgenic mice display exacerbated collagen induced arthritis and bleomycin induced skin fibrosis driven by Type 1 IFN signaling^49,50^ and sex difference in autoimmunity can be driven by Type 1 IFN. Thus, we tested if ZIA was exacerbated in huTLR8 mice using both full dose and half dose zymosan in both male and female mice but did not observe an effect of the transgene, however, we did also observe sex differences when ZIA regardless of the transgene (**Supplemental Figure 2**). The time course of arthritis in female mice was evaluated and scored by a musculoskeletal pathologist (**Supplemental Figure 3**).

**Figure 1.**
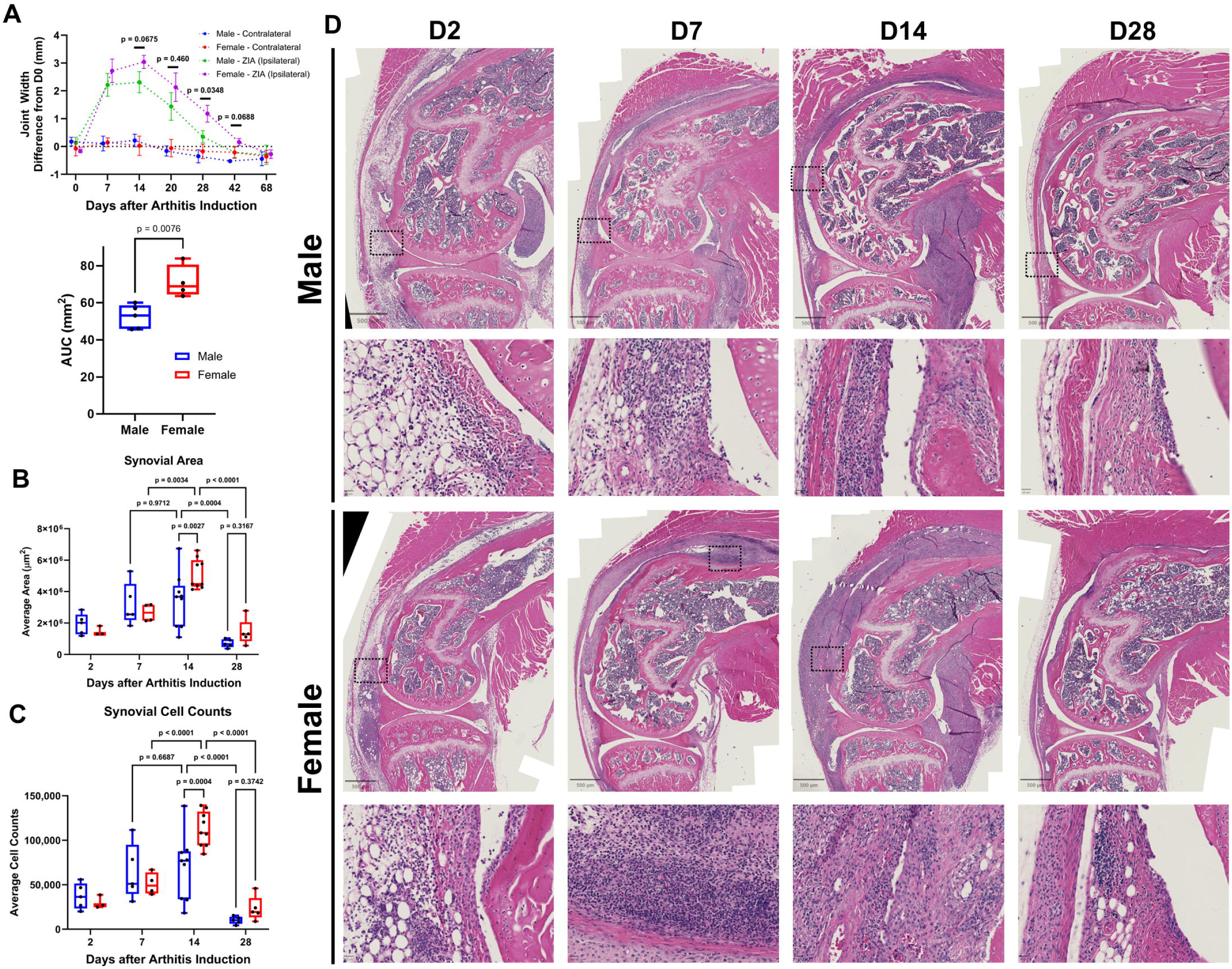
ZIA is a sexually dimorphic, acute, resolving inflammatory arthritis. A) Upper panel. Both ipsilateral (injected with zymosan) and contralateral joint widths were measured every ∼7 days after arthritis induction with 180 µg of Zymosan. Repeated measures ANOVA (Significant main effect of sex), with Tukey’s post-hoc test within day to test each day sex differences. N=4-5 per group, M ± SD. Lower panel. The area under the curve (AUC) was calculated for the male and female ipsilateral joint widths. T-test, n=4-5 per group, each dot represents a mouse, box plots are Min, 25%tile, Mean, 75%tile, and Max. B-D) 180 µg of Zymosan was injected IA in male and female mice. Standard H&E histology was performed to assess pathology at D2, D7, D14, and D28 after arthritis induction. B-C) Synovial tissue area and cell count within that area were measured with the CPath-Arthritis^82^ model. 2-Way ANOVA (Sex vs Day, with a significant main effect of sex) with Tukey’s post-hoc test within day to test each day sex differences. Each dot represents one mouse (average of 3 anatomic levels in the medial compartment), box plots are Min, 25%tile, Mean, 75%tile, and Max, n=4 per group. D) Representative low power (10x, scale bar = 500 µm, upper panels) and high power (100x, scale bar = 20 µm, lower panels) sagittal H&E images of male and female knees at D2, D7, D14 and D28 after arthritis induction. Note the box that identifies the area of high power

Consistent with the CPath-Arthritis results, inflammatory infiltrates peaked at D14 and persisted to D28. Synovial hyperplasia was detected starting on D2; myeloid cells were present throughout the entire disease course, with lymphocytes observed at later time points (Supplemental Figure 3).

We used flow cytometry to test whether differences in synovial infiltration by major immune cell types were associated with sex-based differences in arthritis severity, and performed multiplex ELISA to assess serum levels of cytokines/chemokines. While serum IL-6 and KC (CXCL1, a neutrophil chemokine) were increased at D2 in female mice, no changes in synovial total cell counts, fibroblasts (a potent source of IL-6), neutrophils, monocytes, macrophages, or monocyte-derived DCs were observed between the sexes (**Figure 2**). These results suggest that the increased severity of arthritis in female mice is not related to differences in synovial abundance of major myeloid cell types.

**Figure 2.**
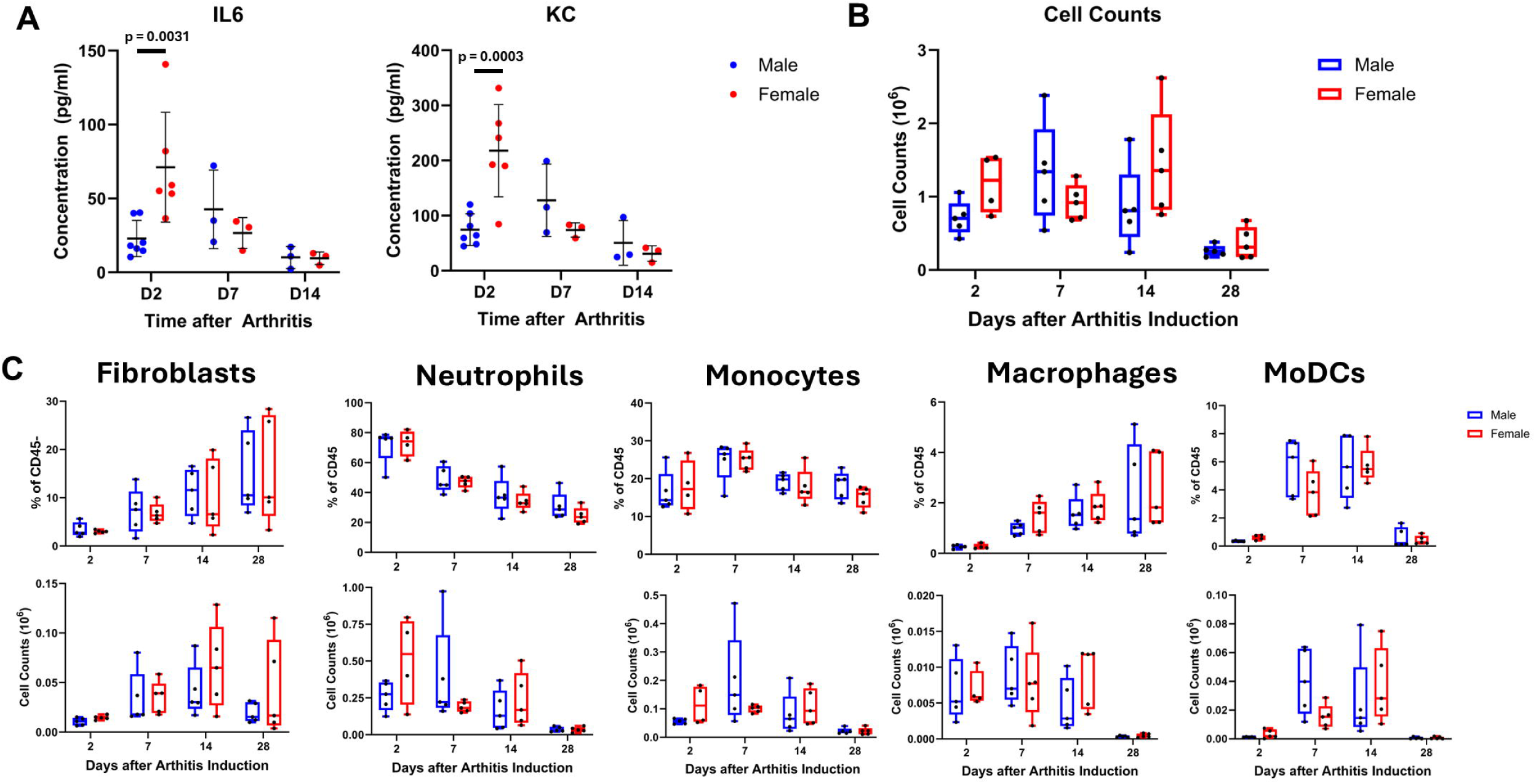
ZIA induces an early sexually dimorphic cytokine and chemokine signature with minimal synovial cell population differences. A) Multiplex serum ELISA of IL-6 and KC (CXCL1) at D2, D7, and D14 after arthritis induction in male and female mice (n=3-7 per group, each dot represents 1 mouse, Mean ± SD). Two-way ANOVA (Significant interaction of Sex and Time) with Tukey’s post-hoc. B) Total cell counts in peri-articular tissue collected for flow cytometry at D2, D7, D14 and D28 after arthritis induction (n=4-5 per group, each dot represents 1 mouse, Box plots are Min, 25%tile, Mean, 75%tile, and Max). C) Percent of CD45+ cells and cell counts of fibroblasts, neutrophils, monocytes macrophages and MoDCs in peri-articular tissue at D2, D7, D14 and D28 after arthritis induction (n=4-5 per group, each dot represents 1 mouse, Box plots are Min, 25%tile, Mean, 75%tile, and Max).

### Distinct synovial myeloid cell subsets in ZIA

Recent work has linked inflammatory arthritis in humans with distinct pathogenic myeloid cell subsets defined by differences in gene expression as determined by single cell RNA sequencing (scRNAseq)^3,4^. To determine whether ZIA recapitulates any of these subsets, we performed scRNAseq on synovial CD45+ immune cells that had been depleted of neutrophils at D2, D7, D14, and D28 of ZIA in male and female mice. Four pooled samples (n=2 female and n=2 male) per timepoint of 3-4 mice per pool with 1 repeat at D7 were collected, resulting in n = 20 scRNAseq samples from ∼80 mice capturing ∼130,000 sequenced high-quality cells. Consistent with the flow cytometry, our scRNAseq analysis revealed that ZIA induces a predominant myeloid infiltration (∼70% of all cells) with smaller numbers of T cells (∼20%), B cells (∼10%) and NK cells/ILCs (**Supplemental Figure 4A**).

We then subclustered the myeloid cells to obtain monocyte and macrophage subsets (**Supplemental Figure 4A)**. After dimensionality reduction, unsupervised clustering and projection onto UMAP space, we observed 13 cell clusters (**Figure 3A**), which we annotated and named using a combination of manual annotation based on cluster marker genes (**Figure 3B**), expression of the top 50 differentially expressed genes (DEGs) in each cluster (**Figure 3C**) and single R^51^(**Supplemental Figure 5A**). Monocytes and macrophages represented the major myeloid population and clustered together, with clusters C1, C3 and C5 classified as monocytes (Mo) based upon expression of *Ly6c2*, *Plac8* and *Ccr2*, cluster C8 as macrophages (Macs) based upon expression of *Adgre1* (encodes F4/80), *Apoe*, *Mrc1* and *C1qa*, and additional clusters of intermediate phenotypes (termed MonoMacs, e.g. C2, C4, C6). Dendritic cells (DCs, C10), monocyte derived DCs (MoDCs, C9) and immature neutrophils (C12, which are Ly6G-low and would not have been removed during cell sorting) were clearly separated from the MonoMac clusters (**Figure 3A**). To distinguish MoDCs from classical DCs, we checked classical monocyte (*Cd14*, *Ly6c2*, and *Ccr2*) and DC markers (*Xcr1*, *Clec9a* and *Batf3*), ensuring both clusters express common DC genes *Dpp4* and *Flt3* (**Supplemental Figure 5B**). Lastly, we calculated cluster purity using neighborPurity (*bluster*, R), which showed that all clusters were well separated (score > 0.85), except for C3, which appeared somewhat diffuse in the center of the UMAP (**Supplemental Figure 5C, Figure 3A**).

**Figure 3.**
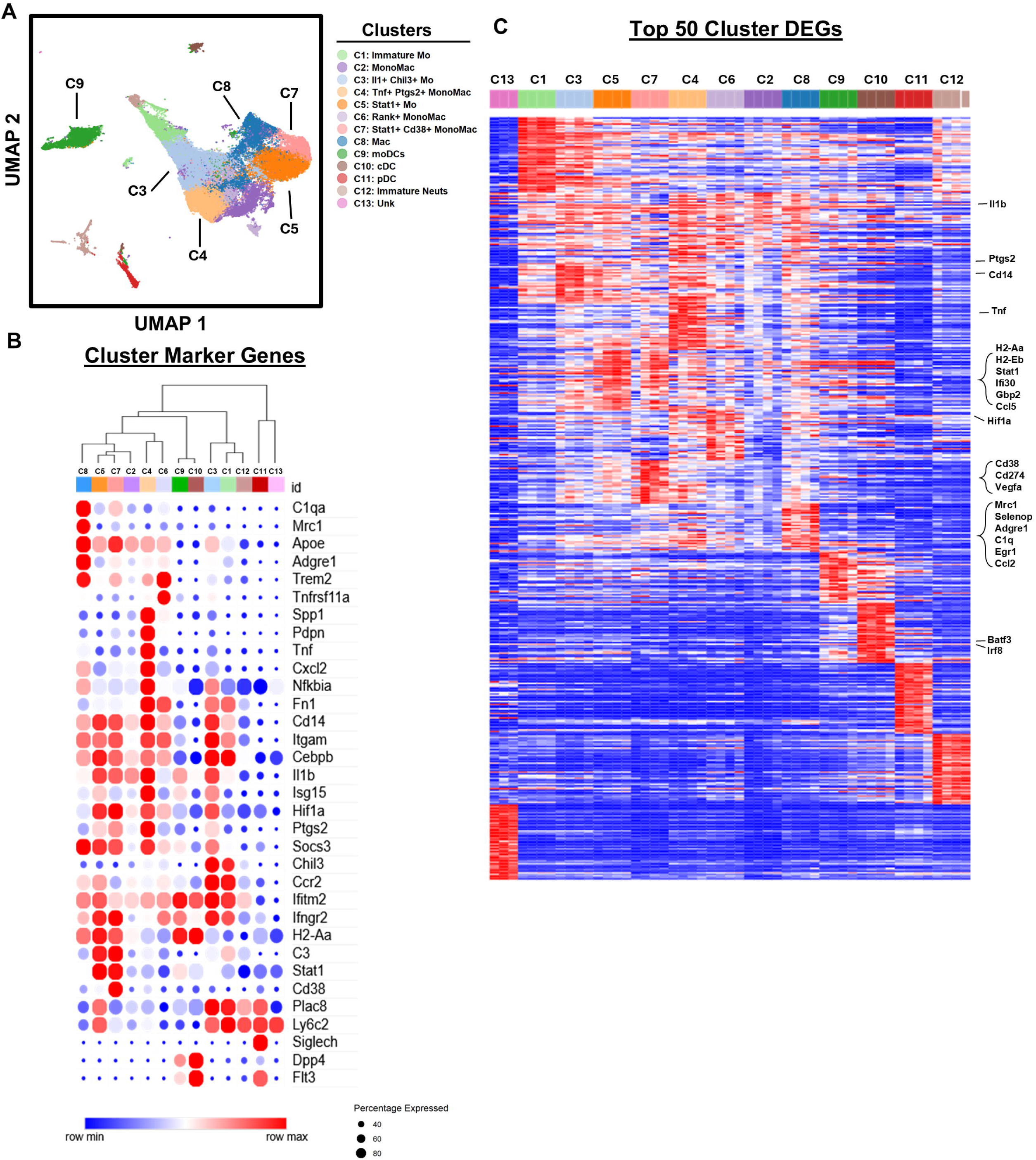
The myeloid transcriptional profile of zymosan induced arthritis. A) Myeloid cells were subclustered and 13 clusters were identified. UMAP representation of the myeloid sub-clustering. B) Curated cluster maker genes that delineate the 13 clusters are shown in a heatmap bubble plot for each myeloid cluster. In addition, hierarchical clustering of the clusters was performed to demonstrate cluster similarity (dendrogram above bubble plot). C) Unique cluster markers were identified with a one vs all strategy (findMarkers, scran) by combining all timepoints and groups to test only between clusters. The top 50 without overlap are shown with hierarchical clustering performed at the cluster level to demonstrate similarity between clusters. Note the select genes on the right side.

Several clusters caught our attention because of high expression of genes implicated in inflammation and RA pathogenesis. Notably, monocyte clusters C3 and C4 expressed various inflammatory NF-kB target genes such as *Il1a, Il1b, Il18, Ccl2, Nfkb1* and *Hif1a*, and select ISGs such as *Ifi204, Ifi207* and *Isg15* that are typically induced by type I IFNs (**Figure 3B-C, Supplemental Figure 5C**). Mo and MonoMac clusters C5 and C7 appeared closely related and co-expressed various inflammatory and IFN target genes such as *Stat1, Cxcl9, Il2rg, Cd274, Ccl5, Ccr5* and various MHC-related genes (**Figure 3B-C, Supplemental Figure 5C**). These results show that ZIA, similar to RA^3,4,52,53^ is associated with subsets of MonoMacs that (co)express inflammatory TNF-NF-kB pathways and an IFN signature. They also raised the question of how closely ZIA MonoMacs and cell clusters model RA inflammatory and pathogenic cells and pathways.

### ZIA recapitulates many of the RA synovial myeloid cell clusters

To evaluate how well ZIA models RA, we undertook three independent and complementary computational approaches to calculate the similarity of ZIA myeloid cells and subsets to an extensive RA synovial single cell myeloid dataset^4^. These methods are: 1) dataset harmonization^54^, 2) reference mapping^55,56^ and 3) cell type prediction using machine learning classification modeling ^57^. Dataset harmonization merged the majority of mouse and human cells into overlapping clusters (**Figure 4A**, left panel). Examination of the original cluster assignments of RA myeloid subsets (**Figure 4A**, middle panel) and ZIA myeloid subsets (**Figure 4A**, right panel) showed similar locations on the UMAP of mouse C3 Chil3+ and human M7 IL1B+ FCN1+ HBEGF+ clusters (blue arrow), mouse C4 TNF+ PTGS2+ and human M4 SPP1+ clusters (red arrow), mouse C7 STAT1+ and human M6 STAT1+ CXCL10+ (pink arrow), mouse C9 moDCs and human M10, M12, and M14 (dendritic cell subsets, green arrow), and mouse C8 Mac with human M1 MERTK+ SELENOP+ LYVE1- clusters (yellow arrow). The relationships of these mouse ZIA and human RA clusters were supported by a more quantitative approach of calculating the Euclidean distance between the centroids of human clusters and their closest mouse cluster (**Figure 4B** and **4C**, key associations noted in the dashed colored ovals). While not our major focus, a good internal control is that both the mouse and human plasmacytoid dendritic cells are nearly 100% overlapping (**Figure 4B** and **4C**).

**Figure 4.**
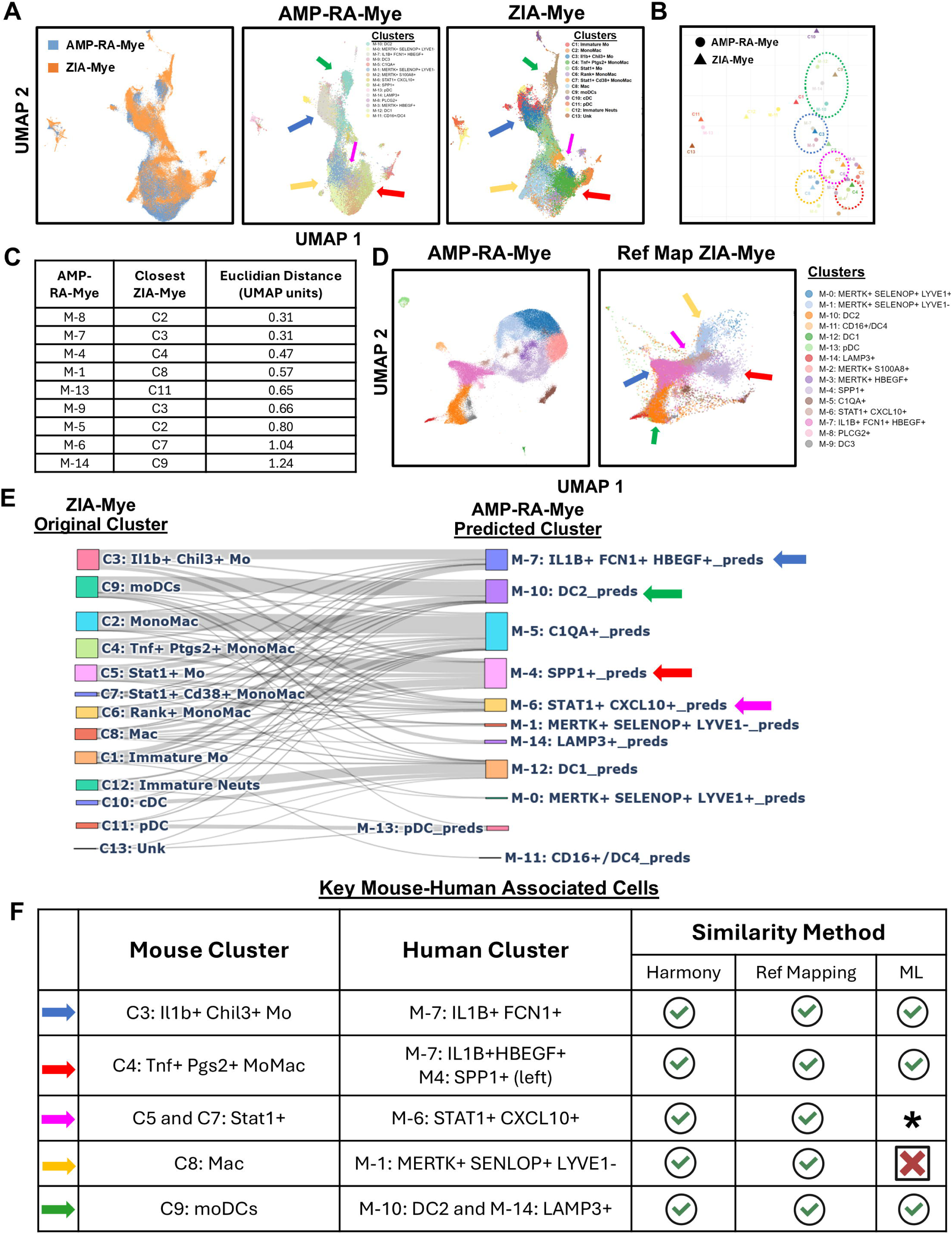
Zymosan induced arthritis mimics RA synovial effector cell types. A) Mouse and human (AMP-RA Phase 2 Myeloid Cells, syn52297840) datasets were merged and harmonized. UMAP plotting of both datasets together (left panel) with AMP-RA-Mye in blue and ZIA-Mye in orange. Each dataset (AMP-RA-Mye = middle panel, ZIA-Mye right panel) is also plotted individually with the original cluster labels. Note the green, blue, yellow, red and pink arrows indicating similar locations of the mouse C9: MoDCs and human M-10: DC2, mouse C3: Il1b+Chil3+ and human M-7: IL1B+ FCN1+ HBEGF+, mouse C8: Macs and human M-1: MERTK+ SELENOP+ LYVE1-, mouse C4: Tnf+Ptgs2+ and human M-4: SPP1+, mouse C7: Stat1+Cd38+ and human C6:STAT1+CXCL10 respectively. B) Centroids of each mouse and human cluster were calculated and ploted with the AMP-RA-Myeloid clusters in circles and the ZIA-Mye in triangles. C) The Euclidean distance from each human cluster to the nearest mouse cluster was calculated and in the table are each of the human clusters with their closest mouse cluster and the distance (in UMAP arbitrary units). The table is ordered by smallest (most similar) to largest distance (cut off at 1.25). D) Reference mapping of ZIA myeloid cells onto predefined RA clusters using PCA anchor point analysis. Left panel. UMAP plot of the myeloid scRNAseq dataset from *Zhang et al., 2023* (syn52297840). Mouse cells were projected into human UMAP space (Right Panel) using the MapQuery function (Seurat) and their human cluster assignments estimated. E) A gradient boosted decision tree model was trained to classify human cell clusters with a nested 5-fold cross validation training strategy and hyperparameter grid search. The most performant (via F1 scores) model was used to infer the human cell cluster of each of the ZIA-Mye cells. After eliminating un-predicted cells and low cell number (<10) predictions, a Sankey diagram demonstrates the flow of cells from their original mouse cell cluster to the predicted human cell cluster. F) A summery table detailing the qualitative success for each of the putative pathogenic clusters of the three similarity methods. A check marks denotes qualitative success, asterisk denotes partial mapping and a X denote failure to map.

Next, we used reference mapping^55,56^ to project the mouse ZIA synovial myeloid cells onto the human RA myeloid cell UMAP space (**Figure 4D** This independent approach assigned a large fraction of the mouse cells into the human M7 IL1B+ FCN1+ HBEGF+ (blue arrow), M-4 SPP1+ (red arrow), M6 STAT1+ CXCL10+ (pink arrow) and M-1 MERTK+ SELENOP+ LYVE1- (yellow arrow) cluster similarities that were identified by the dataset harmonization approach described above. Additionally, a substantial number of mouse cells mapped onto the M-10 DC2 (green arrow) human cluster (**Figure 4D**). Collectively, the data supports a correspondence between at least 4 ZIA and RA myeloid cell clusters, and further suggest that additional ZIA cells annotated as moDCs that did not cluster together with MonoMacs in our scRNAseq analysis are most closely related to RA M10 DC2 cells.

Lastly, we use a machine learning approach to build a cell cluster classifier^57^ (Gradient Boosted Decision Tree) from the human normalized log counts. This allows us to use completely distinct computational methods, i.e. tree-based predictions vs dimensionality reduction-based methods, to assess cell similarity between the mouse and human datasets. The difficulty of training a 15-class model (trained on the 15 RA clusters) is shown in **Supplemental Figure 6** in which we achieved very low performance from 4 cell types (M-9: DC3, M-3: MERTK+ HBEGF+, M-8: PLCG2+, M-6: STAT1+ CXCL10+). However, the remaining cell cluster predictive performance was between 0.76-0.97 F1 with the overall class frequency weighted F1 = 0.78 ± 0.01 suggesting a modestly well performing model. 57% of ZIA cells predicted into RA clusters, which is also indicative of a reasonably performing model, especially when taking into consideration differences between mouse and human genomes, and consistent with other work using a similar approach^58^. We next asked the model to predict which human clusters the mouse cells would fit into (**Supplemental Figure 6** and **Figure 4E**). Notably, large numbers of ZIA cells predicted into the M7 IL1B+ FCN1+ HBEGF+, M4 SPP1+, M6 STAT1+ CXCL10+ and M10 DC2 clusters (**Supplemental Figure 6**). When taking into account the ZIA cluster that cells originated from, this analysis showed more diversity in cell type assignments than the other two methods, but overall the results were largely the same (**Figure 4E**). Specifically, C3 Il1b+Chill3+ monocytes were predicted into the M-7: IL1B+ FCN1+ HBEGF+ (blue arrow), the C4 TNF+ Mo into the M-4: SPP1+ (red arrow), and the MoDCs into the M-10: DC2 human clusters (green arrow). Lastly, despite coming from different ZIA cluster sources many ZIA cells were predicted into the RA STAT1+ CXCL10+ cluster (pink arrow). Overall, the results demonstrate that ZIA myeloid cells and clusters closely recapitulate several RA myeloid cell clusters that have been proposed to be pathogenic based upon expression of inflammatory and IFN pathways (**Figure 4F**).

### ZIA myeloid cells recapitulate ICI-arthritis synovial cells and express a (TNF + PGE2) signature

Severe inflammatory arthritis develops in approximately 4% of cancer patients treated with immune checkpoint inhibitors^6,59,60^. Similar to RA, immune checkpoint inhibitor arthritis (ICI-A) is characterized by infiltration of synovium by immune cells, including IL-1b+ monocytes and IFN-g-expressing T cells^61^. We recently showed that ICI-A synovial myeloid cells map primarily onto 4 RA synovial myeloid clusters: M7 IL1B+ FCN1+ HBEGF+, M4 SPP1+, M6 STAT1+ CXCL10+, and M3 MERTK+ HBEGF+, with smaller numbers of cells mapping onto M1 MERTK+ SELENOP+ LIVE1- and M10 DC2 clusters^14^. Given that five out of these six RA clusters shared by ICI-A were modeled by ZIA cells, we directly tested the relationships of ZIA and ICI-A cells using reference mapping and cell type prediction using machine learning classification modeling **(Figure 5A and B)**. ZIA cells mapped primarily to the ICI-A Cluster 0, Cluster 4, and Cluster 5 with some overlap in Cluster 1 and Cluster 2 (**Figure 5A**). The cell classification modeling performance was excellent as 67% of ZIA cells predicted into ICI-A clusters, with F1 scores between 0.89 – 0.97 (**Figure 5B**, Table). The largest number of ZIA cells predicted into ICI-A Cluster 4 (**Figure 5B**), which we previously showed expresses genes synergistically induced by TNF and PGE2, termed TP genes^8^. Interestingly, ICI-A Cluster 4 was the cluster into which cells from ZIA C3 Chil3+, C4 Tnf+ Ptgs2+ and C5 Stat1+ clusters predicted (**Figure 5B**). These results show effective modeling of ICI-A synovial myeloid cells by ZIA myeloid cells, and highlight the correspondence of three RA-relevant and inflammatory ZIA clusters with ICI-A Cluster 4.

**Figure 5.**
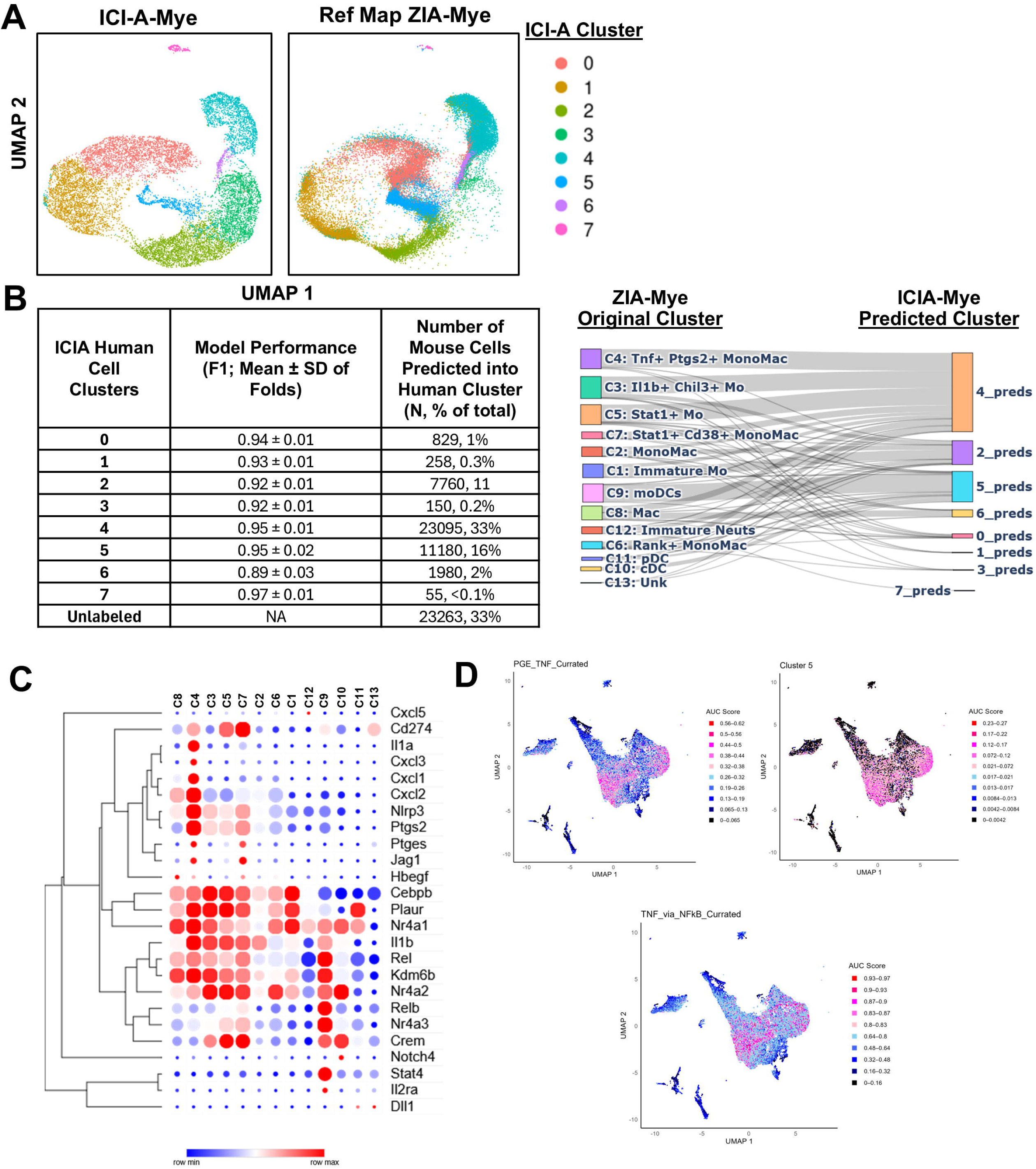
Zymosan induced arthritis maps to PGE+TNF+ immune checkpoint inhibition induced arthritis myeloid cluster. **A)** Reference mapping of ZIA myeloid cells onto predefined ICI-A clusters using PCA anchor point analysis. Left panel. UMAP plot of the myeloid scRNAseq dataset from *Sohki et al 2025* Figure 2E. ZIA-Myelloid cells were projected into human UMAP space (Right Panel) using the MapQuery function (Seurat) and their human cluster assignments estimated. **B)** A gradient boosted decision tree model was trained to classify human cell clusters with a nested 5-fold cross validation training strategy and hyperparameter grid search. The performance of the models is presented in the left table and he most performant (via F1 scores) model was used to infer the human cell cluster of each of the ZIA-Mye cells. After eliminating un-predicted cells and low cell number (<10) predictions, a Sankey diagram demonstrates the flow of cells from their original mouse cell cluster to the predicted human cell cluster. **C)** ZIA-Mye cell clusters plotted with PGE-TNF pathway genes. **D)** AUCell was used to calculate the per cell pathway scores of PGE-TNF curated list, the cluster 5 gene list from *Sohki et al 2025* Figure 1C, and the Hallmark TNF via NFkB list.

Genes synergistically induced by TNF+PGE2 constitute a TP signature that includes genes in pathogenic IL-1, Notch, neutrophil chemokine and Jak-STAT pathways, and is expressed in RA and ICI-arthritis^8,14^. In line with these results, canonical pathogenic TP genes were expressed in ZIA myeloid cell clusters (**Figure 5C).** We further investigated the expression of TP genes in ZIA myeloid clusters by calculating gene set activity scores in each cell using the AUCell package^62^ as described in ‘Materials and Methods’, and visualizing gene expression on UMAP plots. Three TP gene lists were used to identify the distribution of TP-expressing cells, a curated TP list, and the top 20 genes from the two TP clusters in Sokhi et al^14^ (termed TP Cluster 5 and TP Cluster 1). The Hallmark TNF signaling via NF-kB gene list was used as an ‘control’ TNF gene list to identify the general NF-kB response. Expression of TP genes was most highly enriched in the C4 TNF+ PTGS2+ ZIA cluster (**Figure 5D**, top two panels)), which aligns with TP gene expression in the corresponding M7 IL1B+ FCN1+ HBEGF+ RA cluster^14^ . The TP signature was also apparent in a subset of C8 Macs and in adjacent areas of subsets of C5 STAT1+ and C7 STAT1+ CD38+ clusters. This distribution of the TP signature across more than one ZIA cluster was also observed in RA^14^. In contrast to TP genes, genes in the TNF signaling via NF-kB pathway were more broadly expressed and were enriched in C3, suggesting a lack of PGE2 signal in this cluster (**Figure 5D**, bottom panel). These results suggest that ZIA recapitulates the interplay between TNF and PGE2 signaling that occurs in subsets of RA and ICI-A myeloid cells.

### Temporal evolution of distinct myeloid cell subtypes infiltrating the synovium in ZIA

We reasoned that analysis of the temporal evolution of synovial myeloid subsets and their pattern of gene expression would yield insights into mechanisms that drive the induction, maintenance and resolution phases of an episode or flare of inflammatory arthritis. The early phase of ZIA arthritis (D2) was dominated by RA-relevant clusters C3 Il1b+ Chil3+, C4 Tnf+ Ptgs2+ and C9 MoDCs (**Figure 6A and 6B**). These cell populations were associated with the ramping-up phase of arthritis, and their percent populations progressively decreased to D14. Gene set enrichment analysis showed that, on D2, C3 Il1b+ Chil3+ and C4 Tnf+ Ptgs2+ cells showed most significant enrichment of ‘TNF signaling via NF-kB’ and inflammatory pathways, whereas C9 MoDCs co-expressed NF-kB and IFN pathway genes (**Supplemental Figure 6**). Pathway scoring clearly demonstrates the greater magnitude of enrichment TNF signaling relative to IFN signaling in C3 and C4 on D2, when these are the major clusters in the inflamed synovium (**Figure 6C**). These results indicate that the induction phase of arthritis was associated predominantly with NF-kB signaling, with a smaller population of C9 moDCs co-expressing inflammatory and IFN pathways even at early stages of arthritis.

**Figure 6.**
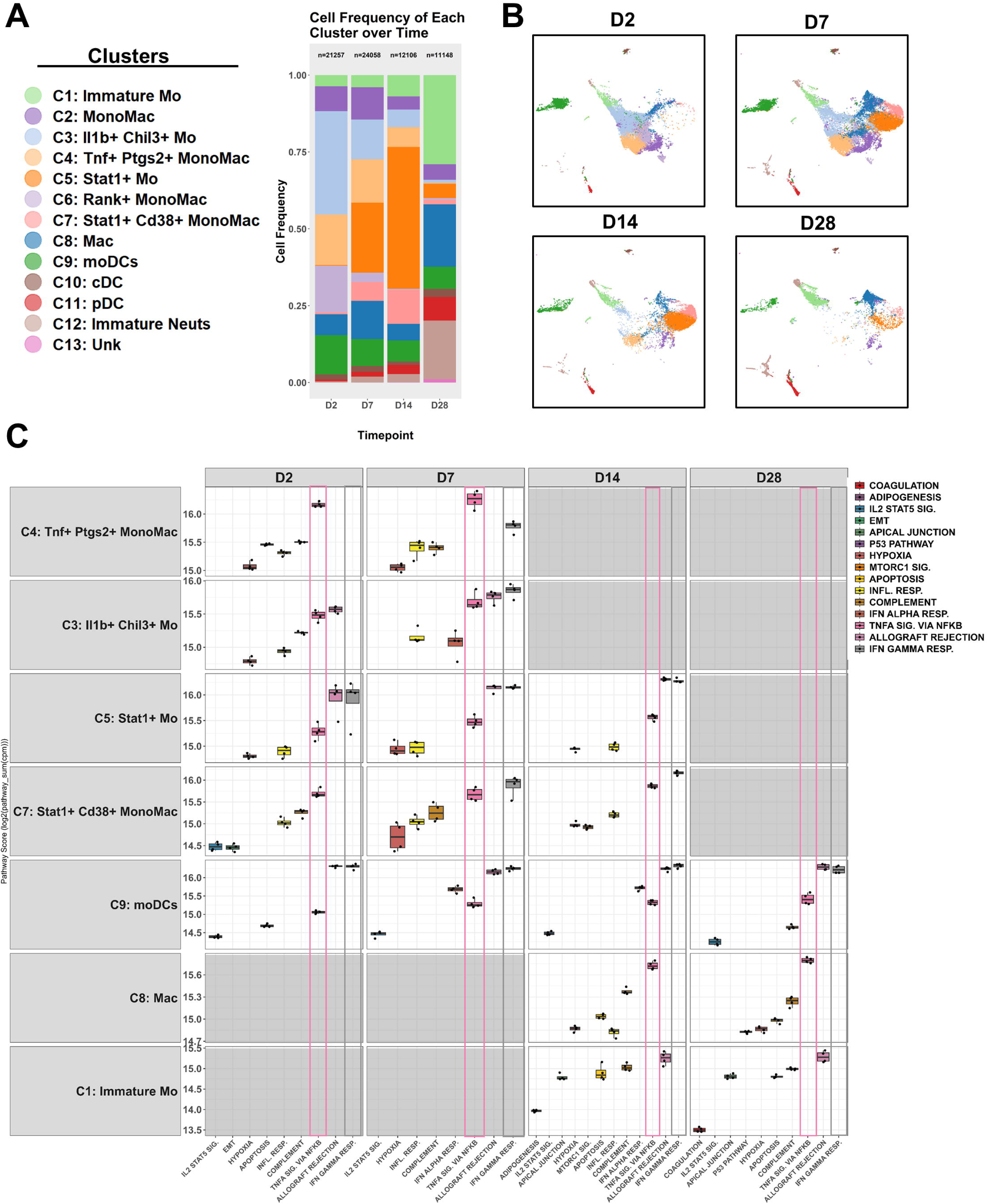
The myeloid temporal and pathway profile of zymosan induced arthritis. A) Cluster frequency at D2, D7, D14 and D28 of the ZIA synovial myeloid cells. B) UMAP plots at D2, D7, D14 and D28 demonstrating the distribution of cells over time. C) Psuedo bulk analysis at each timepoint for each cluster and differential gene analysis (one vs all) between clusters was performed. Pathway analysis from the DEGs was performed and the top 5 significant pathway scores of putative effector cells during each phase of disease is plotted. Clusters whose cell frequency falls below 10% in a timepoint were not considered for analysis. Data are presented as boxplots, where each point is the log_2_ sum of all pathway genes pseudo bulk CPM (n=4 per group). Note the TNF_via_NFkB (Pink) and IFN gamma (Grey) pathways highlighted.

The peak of ZIA at D7-D14 was marked by the appearance of large numbers of the related C5 Stat1+ and C7 Stat1+ Cd38+ cells on D7, both of which together expanded to form the predominant myeloid cells population on D14 (**Figure 6A and 6B**), suggesting that these are the effector cells during the middle phase of arthritis. Interestingly, C5 and C7 cells co-expressed genes in the NF-kB and IFN pathways, with higher expression of IFN pathway genes. These clusters also highly significantly expressed genes in the ‘Allograft Rejection’ pathway (this gene set is comprised of a mix of inflammatory genes and ISGs) **(Supplemental Figure 6** and **Figure 6C**). The emergence of an IFN signature in C5 and C7 cells was associated by infiltration of synovium by IFN-g-expressing T cells and NK cells (**Supplemental Figures 4A, 7A and 7B**). Expressed *Ifng* most likely reflects cytokine-mediated activation of NK cells and bystander activation of T cells, and provides an explanation for the observed IFN signature.

At D28, when arthritis was resolving, the above-described cell clusters had mostly resolved, and the synovium was predominantly populated by C1 immature Mo, C8 differentiated macrophages, and C12 immature neutrophils. Instead of inflammatory pathways, the C1 immature Mo were most significantly enriched in genes in the ‘complement’ and ‘apical junction’ pathways. In contrast and in accord with ongoing inflammation as observed on histology (**Figure 1**), the C8 macrophages continued to express IFN-g and NF-kB pathways (**Figure 6C**). These results clearly link specific myeloid subsets with distinct phases of arthritis and suggest that cells driven by NF-kB signaling predominate at early phases of disease, followed by emergence of cell clusters that co-express IFN-Jak-STAT and NF-kB pathways.

### Sexually dimorphic cellular populations and gene signatures

To address the question of whether any of these myeloid cell clusters contribute to the sexually dimorphic phenotype we performed both cell abundance analysis and pseudo bulk differential gene expression and pathway analysis between the sexes within each cluster. UMAP contour plots and cell frequency analysis demonstrates that both C5 and C7 were increased in the female samples (**Figure 7A**). Interrogating cell frequency showed that female samples have nearly 3x the number of C5 Stat1+ cells (Male ∼10% vs Female ∼30%) and C7 Stat1+ Cd38+ cells (Male ∼2.5% vs Female ∼7.5%) at D7 (**Figure 7B**). In addition to the higher abundance, when we performed pseudo-bulk analysis and calculated count normalized (cpm) pathway scores in these two clusters, female C7 cells showed higher pathway scores in the metabolic and activation-related pathways oxidative phosphorylation, Mtorc1, and glycolysis relative to male C7 cells (**Supplemental Figure 8A** and **Figure 7C**). Thus, on D7, females exhibit higher numbers of ISG^hi^ C5 and C7 cells that are metabolically active. In contrast to C5 and C7 that peak on D14, the frequency of NF-kB-predominant IL1B+ CHIL3+ C3 and TNF+ PTGS2+ C4 cells that peaked on D2 was comparable in male and female mice. However, analysis of gene expression revealed that D2 C3 and C4 cells had strongly enriched IFN signatures in female compared to male cells (**Figure 7D**). To corroborate the early, higher IFN response in female cells, we sorted Ly6C+, CD11b+ synovial myeloid cells on D2 and performed qPCR analysis of ISGs *Ifit1*, *Cxcl10*, *Isg15* and *Mx1*, which were all higher in female mice (**Supplemental Figure 8A**), while classical NF-kB targets *Il1β*, *Tnf*, and *Il6* were not different between the sexes (**Supplemental Figure 8B**). Overall, these results link an increased IFN response, as evidenced by significantly higher ISG expression on D2 and a 3-fold increased in ISG-expressing Stat1+ C5 and C7 clusters on D7, to increased arthritis in female mice.

**Figure 7.**
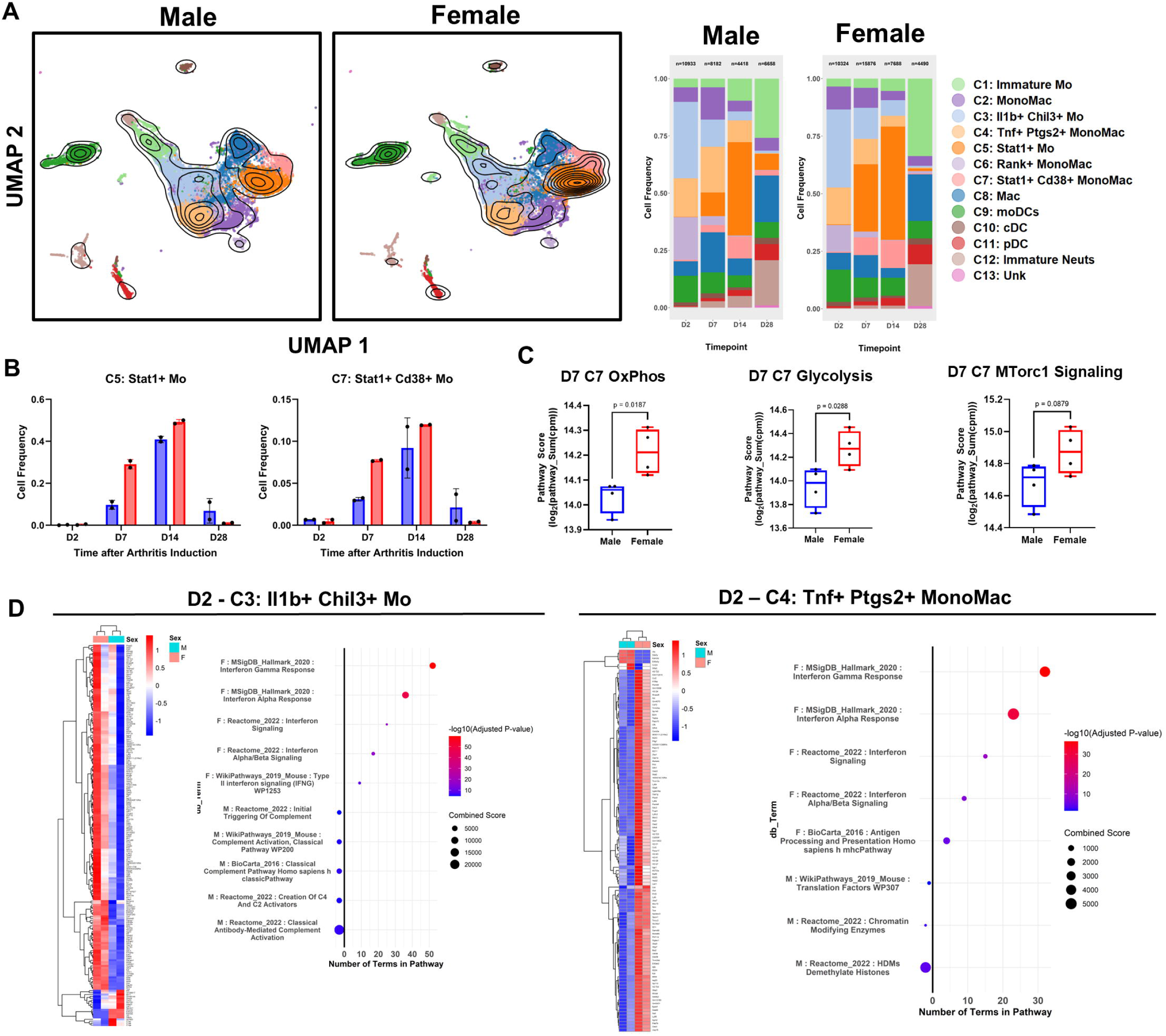
Sexually dimorphic cell cluster abundance and transcriptional pathways in ZIA. A) Contour UMAP analysis of male and female samples at all timepoints (left two panels) and cell frequency of each cluster at each timepoint between the sexes (right two panels). B) Cell frequencies of Cluster 5 : Stat1+ Mo (Left) and Cluster 7 : Stat1+CD38+ Mo (Right) of each male (Blue) and female (Red) sample at D2, D7, D14, and D28 after arthritis induction. C) Psuedo bulk, DEG and cluster analysis between sex was performed at D7. Significant Hallmark cell metabolism pathways (Oxidative Phosphorylation, Glycolysis, and MTorc1 Signaling) are plotted. Data are presented as boxplots, where each point is the log_2_ sum of all pathway genes pseudo bulk CPM (n=4) D) Psuedo bulk, DEG and cluster analysis between sex was performed at D2, D14 and D28 (n=2 per group). DEG and significant pathways for D2: C3 - Il1b+Chil3+ Mo and D2 : C4 – Tnf+Ptgs2+MonoMac are shown.

## Discussion

Important challenges in the RA field are to understand the genesis and pathogenic role of newly identified subsets of immune cells that infiltrate inflamed synovium, and the sexually dimorphic nature of disease. In this study, we have utilized complementary computational strategies to rigorously show that the ZIA model effectively recapitulates myeloid cell subsets previously identified in RA and ICI-A, and are proposed to be pathogenic based upon expression of inflammatory genes. These shared RA, ICI-A and ZIA synovial myeloid cells express genes in NF-kB, IFN-STAT and PGE2 pathways. In ZIA, NF-kB and PGE2 pathways predominate and interact during the initial induction phase of arthritis, followed by a sustained phase characterized by emergence of MonoMac subsets that co-express IFN and NF-kB pathways. This work establishes ZIA as a relevant model for RA and ICI-A that enables dissection of key aspects of pathogenesis of the effector phase of inflammatory arthritis. We have taken the first step towards this goal by showing that ZIA is sexually dimorphic, and linking increased arthritis in female mice with early phase induction of IFN signaling in NF-kB-predominant MonoMac subsets, and strongly increased infiltration by myeloid cell subsets that co-express ISGs and NF-kB target genes during the sustained phase of disease.

Although the application of high dimensional single cell profiling coupled with in depth computational analysis has transformed our understanding of the immune landscape of RA, there is little information about how well various arthritis models recapitulate the phenotypes of newly discovered cell subsets that have been proposed to be pathogenic in patients. Previous work using scRNAseq analysis of mouse arthritis was performed solely in the serum transfer-induced arthritis model that does not recapitulate the RA/ICI-A synovial IFN signature, comparison of mouse to human cells was limited in scope^7,63^. Our study advances the field by capturing the IFN signature and developing a comprehensive computational approach to rigorously identifying mouse arthritis myeloid cells and subsets that closely match their human disease counterparts. The approach involves using the find orthologs function (*gorth*, gprofileR, R) to assign identity between mouse and human genes followed by utilization of 3 distinct but complementary computational methods to relate mouse and human disease myeloid cells and subsets. It is reassuring that the 3 different approaches yielded similar results, and by focusing on mouse-human cell relationships commonly identified by all 3 methods we have minimized false positives and identified core mouse myeloid cells and subsets that accurately model human disease cells and subsets. We propose that application of this strategy to various mouse arthritis models will identify which aspects of RA are best recapitulated by which model. This will in turn inform and enhance the relevance of mechanistic investigation and therapeutic testing in mouse models.

The application of this computational strategy to ZIA clearly identified two related ZIA myeloid cell clusters, C3 Il1b+ Chil3+ and C4 Tnf+ Ptgs2+, that closely model two RA cell clusters M7 IL1B+ HBEGF+ and M4 SPP1+ (‘left subset’) that similarly are adjacent to each other in UMAP space. Both of these clusters, in RA and ZIA, express inflammatory target genes such as *IL1B*, and the overlap region between M7 and the left subset of M4 is notable for expression of genes synergistically induced by TNF and PGE2 (termed TP genes)^14^ that include neutrophil chemokines such as CXCL2 and EGFR ligands such as HBEGF that promote fibroblast invasiveness, potentially linking these clusters to tissue destructive disease.

Interestingly, this overlap region on the RA UMAP also corresponds to a myeloid cell neighborhood found in the myeloid subtype of RA^4^. The ZIA C4 cluster is interesting from a pathogenic point of view as it is the only cluster expressing *TNF*. The identification of C3/C4 myeloid subsets sets the stage for future work to identify molecular pathways that induce these cells and drive pathogenic gene expression, and mechanisms by which these myeloid cells contribute to disease. Of particular relevance for RA pathogenesis will be to identify what drives *TNF* expression in C4, and the role of PGE2 pathways in tissue destructive disease. ZIA C9, which we annotated as MoDCs, also closely corresponds to an RA cluster, namely M10, annotated as DC2 in ref 8. The ZIA C9 and RA M10 clusters share an IFN signature related to MHC gene expression, and RA DC2s colocalize with T cells in the sublining layer^13^. It is tempting to speculate that ZIA moDCs may similarly colocalize with T cells and may play a role in the T cell activation and IFN-g expression that we observed in this model.

ZIA exhibited two interesting myeloid subsets, C5 Stat1+ and C7 Stat1+ Cd38+ cells, that co-expressed inflammatory NF-kB targets and ISGs, and thus recapitulate the co-activation of these pathways in myeloid cells not only in RA and ICI-A, but also in various inflammatory conditions such as lupus nephritis, inflammatory bowel disease and COVID-19 lungs^3,4,15,64,65^.

Although the ZIA C7 cluster to a limited extent recapitulated the RA M6 STAT1+ CXCL10+ cluster, overall, the C5 and C7 clusters did not well correspond to specific RA clusters. Instead, the individual cells contained within C5 and C7 clusters distributed amongst three pathogenic RA clusters (M4, M7 and M10), in addition to M6, by both representation mapping and the computationally rigorous machine learning approaches. Thus, these cells are highly relevant for RA pathogenesis; most likely the mapping of these cells into distinct RA clusters reflects differences in the relative strength of NF-kB versus IFN gene expression programs. The mapping of ZIA C5 and C7 clusters onto ICI-A cluster 4 further supports the pathophysiological relevance of these myeloid subsets. Future investigation of these cells will provide insights into the role of IFN signaling, and the interplay between NF-kB and IFN pathways, in arthritis pathogenesis.

Our results suggest ZIA is comprised of two overlapping phases, with an early TNF, PGE2 and NF-kB-driven induction phase followed by a switch to distinct myeloid cell subsets that co-express NF-kB and IFN responses and are associated with sustained arthritis beyond the D7 time point. Human RA, in the absence of effective treatment, is characterized by sustained disease with superimposed flares and spontaneous, albeit partial, remissions that result in a worsening of disease over time. Both RA flares and the induction phase of ZIA are characterized by influx of neutrophils, and one interesting possibility is that RA flares are induced by signals that induce C3 and C4 cells and the TP signature. In contrast, sustained disease and the plateau at the peak of each flare may be mediated by the distinct C5- and C7-myeloid subsets that express an IFN signature. This has therapeutic implications for targeting different cells subsets at different phases of disease. As the ZIA-modeled RA M4, M6, M7 and M10 clusters are differentially associated with different subtypes of RA (termed CTAPs^4^), it is possible that investigation of the effects of therapeutics on the corresponding cell clusters in ZIA will yield insights useful for selective targeting of patients with distinct CTAPs. An intriguing possibility is targeting IFN-g, especially as the neutralizing IFN-g antibody emapalumab is FDA-approved for other indications. However, as IFN-g can have homeostatic and suppressive effects upon the stromal and vascular compartments^66^, the role of IFN-g will need to be carefully dissected in animal models like ZIA prior to considering a clinical trial of IFN-g blockade in RA.

The basis for the sexual dimorphism of RA is not well understood. Although the increased incidence of RA in females is the best established sexually dimorphic clinical finding^24–26^, females with RA are more refractory to therapy, less likely to achieve remission, and trend towards higher disease activity than men^27–29^. Our findings linking increased ZIA with an increased IFN response suggest an explanation for increased arthritis in females. It is well understood that the female sex confers a stronger infection response driven by IFN, exemplified by lower infections in the first six months of life and a higher antibody titers after vaccination^67–69^. The type 1 IFN response in lupus, one of the most sexually dimorphic autoimmune diseases, is also well documented^26^ and the sex differences are, at least in part, driven by overexpression of nucleic acid sensors on the X-chromosome^70^. However, it is unlikely that aberrant nucleic acid sensing plays a role here in the sex differences as we saw no exacerbation of the phenotype in the Tg8 experiments (Supplemental Figure 2). Further, there is evidence of increased IFN (largely type 1 IFN) activity in the female myeloid compartment at baseline and in stimulated conditions^34,35^. Despite this literature and our results, the role of type 1 vs type 2 IFN in RA and how IFN may interact with sex to alter disease progression is not well understood. It is possible that this increased IFN response contributes to resistance to therapy and lower rates of remission, for example, by epigenetic mechanisms that can induce resistance to inhibitors of inflammatory pathways^71^. One broad challenge of animal models of sexually dimorphic immune-mediated diseases is that they tend to exhibit increased severity rather than increased incidence. It is possible that the increased IFN response we have described can push individuals past a threshold beyond which clinically penetrant disease is observed, but this notion would be need to be tested in more complex models that include an autoimmune component. A previous report shows that female myeloid cells are more responsive to IFN stimulation in vitro, making it likely that the increased IFN response we observed is related to increased responsiveness of myeloid cells to low levels of synovial IFNs. Such an increased cell-intrinsic responsiveness to IFNs may also explain the striking 3-fold expansion of C3 STAT1+ and C7 STAT1+ CD38+ myeloid subsets in female relative to male mice.

There are several limitations to our study. As holds true for all animal models, ZIA only recapitulates select aspects of RA pathogenesis. IFN-g and type I IFNs induce overlapping sets of ISGs, and we have not distinguished between the role(s) of distinct types of IFNs in generating the IFN response in ZIA. Our study raises the interesting possibilities that differential sensitivity to IFNs and expansion of human counterparts of ZIA C5 STAT1+ and C7 STAT1+ CD38+ myeloid cells also contributes to sexual dimorphism of RA in humans. Initial efforts to test this idea using the AMP data set^4^ have not to date detected sex differences, but are limited by relatively small sample size, heterogeneity of patients across 7 different CTAPs (only some of which may be sexually dimorphic), and the distribution of ZIA C5 and C7 cells amongst 4 RA subsets, making a direct comparison of expansion of corresponding cell clusters not feasible.

Our study motivates future analysis of sexually dimorphic myeloid cell subsets and the molecular pathways they express in larger RA data sets, and the development of novel computational strategies to analyze sexual dimorphism of the human cell counterparts of ZIA C5 and C7, either after re-clustering of RA cells or independently of cell cluster.

## Materials and Methods

### Mouse Model of Arthritis

All mouse experiments were performed with prior approval from Weil Cornell Institutional Animal Use and Care Committee. Female and male C57BL/6 mice were purchased from Jackson Labs at 8-10 weeks old and allowed to acclimate to our facility for at least 2 weeks. Zymosan purified from *Saccharomyces A* was dissolved in sterile PBS at 30 mg/ml and boiled for 30 mins. Intra-articular injections of 90, 135 or 180 ug with a 30g needle were performed to induce arthritis. TLR8-Transgenic mice were kindly provided by Dr. Frank Barrat and were originally generated via BAC/ES technologies in the original work^49^.

### Histology

Joints were dissected *en bloc*, incubated on a rocker in 10% neutral buffered formalin for 8-12 hours at room temperature and then incubated in 14% EDTA (buffered with acidic acid) for 14 days at room temperature on a rocker with 2 solution changes. The joints were then embedded in wax and the medial compartment section in the sagittal plain. Three levels separated by at least 100 microns were collected, stained with H&E and slide scanned with a Zeiss Axioscan 7 at 20x. Standard histology scoring^72^ was performed on a subset of images while computational pathology (CPath-Arthritis)^48^ to measure synovial area and cell counts was performed on all slides.

### Joint Measurement

Calipers were used to measure the medial to lateral width of the knee joint at the level of the patella in full extension once a week.

### Serum Multiplex Elisa

At sacrifice, whole blood was collected, allowed to coagulate and spun at 5000g for 20 mins to separate the serum. Serum analyte concentration were measured with the Milipore 36-plex T-cell mouse high sensitivity kit.

### Flow Cytometry

Peri-articular tissue was dissected from the anterior compartment of the injected joints in cold sterile RPMI. Tissue was manually dissociated in the RPMI with a razor blade and then incubated with Dispase (0.8 mg/ml), Collagenase P (0.2 mg/ml) and DNASE I (0.1 mg/ml) at 37C for 45 minutes on a rocker. Tissue preparations were then pipetted up and down 40 times in a long tipped pasture pipette, incubated with EDTA 0.5M for 5 minutes at 37C on a rocker and then passed through a 70 µm strainer. Cell suspension were blocked with Fc block, and then stained with antibody cocktail detailed below for 10 minutes, washed with FACS buffer, stained with DAPI immediately before analysis and flowed on a Symphony A3 (BD). Anti-NK1.1-BUV395, Anti-CD3-BUV563, Anti-Cd11c-BUV737, Anti-Ly6G-BUV805, Anti-CD11b-BV421, Anti-CD26-BV605, Anti-CD4-BV650, Anti-MHCII-BV711, Anti-CD8-BV786, Anti-Cx3cr1-AF488, Anti-Ly6c-RB705, Anti CD90.2-RB780, Anti-B220-PE, Anti-CD9PE/Dazzle594, Anti F4/80-PE/Cy7,Anti-CD31-APC, and Anti-CD45-APC-R700. Flow gating is detailed in **Supplemental Figure 9**.

### Single Cell RNA Sequencing and Analysis

Single cell suspensions were generated and stained as described above in the Cytometry methods. CD45+Ly6G- population from female and male WT and Tg8 mice at D2, D7, D14, and D28 (n=3-4 per group) were sorted, counted and then evenly pooled into scRNAseq reactions on the 10x controller. Mice from this experiment were injected on separate days and euthanized on the same day to minimize batch effect from the tissue disassociation, flow sorting and 10X controller procedures. Subsequently, a repeat D7 experiment was performed with female and male WT and Tg8 mice (n=3-4), sorted and pooled in the same way. This resulted in 20 scRNAseq reactions pooled from ∼80 mice in 2 batches. RNA libraries were generated per the manufacturers recommendations and sequenced on an Illumina Nova X. Data were processed with CellRanger to generate feature count matrixes. In general, analysis was performed following the “Orchestrating Single Cell Analysis with Bioconductor”^76^ pipeline in R (4.5.0). After merging and standard quality control filtering for mitochondrial RNA expression (<10%), library size (>500 counts), unique genes (>200 genes) and removal of doublets (consensus via three methods: doublet cluster identification^73^, doublet simulation^74^, doublet classification^75^), we obtained 130,181 high quality cells. In addition, batch harmonization was performed with Harmony^54^, general cell type annotations were obtained form SingleR^51^ and cell type purity was calculated with *neighborPurity* (bluster). Pathway analysis at each timepoint was performed by pseudo-bulking the four samples (WT-M, WT-F, Tg8-M and Tg8-F) for each cluster per timepoint, performing differential gene expression with edgeR (One vs All), and summarizing the significant DEGs from the Hallmark pathways (mSigDB). Data are plotted as log_2_(sum(all_sig_pathway_genes(counts_per_million))). Sex difference analysis was performed by pseudo-bulking the four samples (WT-M, WT-F, Tg8-M and Tg8-F) for each cluster per timepoint, performing differential gene expression with edgeR between the sexes (n=4 per group at D7; n=2 per group at D2, D14, D28). Differential genes were analyzed with EnricheR^77–79^ to assess pathways differences from the Hallmark, Reactome, Biocarta, and WikiPathways databases.

### Flow Sorted QPCR

Single cell suspension from the synovium were generated as described above and flow sorted for CD45+Ly6g-Cd11b+Ly6c+ myeloid cells. RNA was extracted from these cells, cDNA generated and target gene expression was tested with quantitative polymerase chain reaction (QPCR) via the Syber Green system. Relative percent gene expression to a house keeping (GAPDH) are plotted.

### Human RA Dataset and Analysis

The two human datasets (AMP-RA-Mye and ICI-A) were obtained via access instructions from their original work^4,80^. ZIA and the human data were analyzed for similarity in 3 ways, 1) harmonization^54^, 2) reference mapping^55,56^ and 3) machine learning classification^57^. For all approaches datasets after QC filtering were used and mouse genes were converted into their human homologs with the *gorth funtion* (gprofileR, R). For harmonization, datasets were then merged, highly variable genes selected, PCAs calculated and harmonization^54^ was performed to remove the effect of species source. UMAP on the harmonized PCs was performed, the original clusters were overlayed into the new UMAP space, their centroids calculated and the nearest neighbor cluster from the other dataset via Euclidean distance was calculated. For reference mapping^55,56^, datasets after PCA calculations were used, we found anchors on the in the reference PCA space (e.g. human) with *findanchors* (seruat, R), and mutual nearest neighbor was found between the reference and query (mouse) PCs. Mouse cells were then projected into the original human UMAP and cluster space. For machine learning (scikit-learn framework, python), a gradient boosted decision tree (GBDT, xgboost) was trained on the 15 clusters (i.e. classes) using nested 5-fold cross validation with grid search for the GBDT parameters: learning_rate = [0.1, 0.05], colsample_bytree = [0.6, 0.8, 1.0, subsample = [0.5], max_depth = [2, 4, 8], "n_estimators": [200, 400, 800], gamma = [0], min_child_weight = [1]. Highly variable genes were used as features in the training. The best performing model by weighted F1 was selected and used to infer human cluster on the mouse cells. Plotly was used to create the Sankey plots.

## Dataset Availability

Mouse scRNAseq is available at https://gitlab.com/hssgenomics/. Human data is available via their source manuscripts^4,14,80^.

## Code Availability

All scripts (r or python) are available at Dr. Richard Bell’s github (github.com/rdbell) with environment dependencies.

---

**Supplemental Figure 1 associated with Figure 1. ZIA is a titratable phenotype.**

A-C) Zymosan was titrated to 90, 135, and 180 µg per intra-articular (knee) injection and mice were euthanized 2, 7 and 14 days after injection. Standard H&E histology of knee joints was performed to assess pathology.

A-B) Synovial tissue area and cell count within that area was measured with the CPath-Arthritis model^81^. 2-Way ANOVA (Dose vs Day) with Tukey’s post-hoc test within day to test for the dose effect. Each dot represents one mouse (average of 3 anatomic levels in the medial compartment), box plots are Min, 25%tile, Mean, 75%tile, and Max, n=4 per group.

C) Representative sagittal H&E images of sections from knee joints 14 days after injection of 90, 135, or 180 µg of zymosan.x

**Supplemental Figure 2 associated with Figure 1. Observations of sex differences in Tg8 mice.** Humanized TLR8 (Tg8) transgenic mice display exacerbated collagen induced arthritis and bleomycin induced skin fibrosis driven by Type 1 IFN signaling. We tested if ZIA was exacerbated in huTLR8 mice using both full dose and half dose zymosan in both male and female mice.

A) Tg8 mice injected intraarticularly with 180 µg zymosan have increased serum IFNγ (left panel, T-test n=6 per group), however this was not associated with increased synovial histopathology at either D7 or D28 (Synovial Area = middle panel and Synovial Cell Counts = right panel). Each dot represents one mouse (average of 3 anatomic levels in the medial compartment), box plots are Min, 25%tile, Mean, 75%tile, and Max, n=9-12 per group.

B) To test if there is a ceiling effect at 180 µg, we injected knees with a half dose (90 µg) of zymosan and also saw no difference in histopathology between WT and Tg8 mice at D7(Synovial Area = left panel and Synovial Cell Counts = right panel). Each dot represents one mouse (average of 3 anatomic levels in the medial compartment), box plots are Min, 25%tile, Mean, 75%tile, and Max, n=6-8 per group.

C) We observed various indications that female Tg8 mice had increased pathology, with increased synovial area at D28, more synovial monocytes and NK cells at D2 and more neutrophils at D7. Box plots are Min, 25%tile, Mean, 75%tile, and Max, n=8-12 per group Each dot represents one mouse (Histopathology is the average of 3 anatomic levels in the medial compartment).

**Supplemental Figure 3 associated with Figure 1. Inflammatory infiltrate and pannus noted with fibroblasts, myeloid and lymphoid cells.**

Review and scoring of histopathology by a pathologist.

A) Inflammatory infiltrate scores in female mice at D2, D7, D14 and D28 (n=3) and additional pathology notes.

B) Pannus invasion scores in female mice at D2, D7, D14 and D28 (n=3) and additional pathology notes.

**Supplemental Figure 4 associated with Figure 3. The synovial immune cell landscape in ZIA.**

A) CD45+, Ly6G- peri-articular cells were flow sorted and input into the 10x Genomics single cell RNAseq pipeline. UMAP dimensional reduction and automated cell type (SingleR) analysis is shown in the left panel.

B) Curated marker genes are shown in a heatmap bubble plot for each cell type (Right Panel).

**Supplemental Figure 5 associated with Figure 3. Accessory scRNAseq Myeloid cell clustering information.**

A) Macrophages (Red), Monocytes(Purple), Neutrophils (Brown) and moDCs (Orange) from Supplemental Figure 4 we sub-clustered and automated cell type (SingleR) analysis was performed.

B) Dendritic cell and monocyte marker genes in clusters annotated as monocyte DCs and conventional DCs.

C) Cluster purity scores generated by *neighborPurity* (bluster) for the myeloid sub clustering.

D) Additional gene markers for PGE, TNF and IFN pathways in the myeloid clusters.

**Supplemental Figure 6 associated with Figure 4. Gradient boosted decision tree performance and cell predictions.** A table with each human cluster the models internal performance (F1, mean and standard deviation of folds). In addition, the total number of mouse ZIA-Mye cells predicted into each cluster.

**Supplemental Figure 7 associated with figure 6. Gene set enrichment analysis of synovial myeloid cells in zymosan induced arthritis**. Bubble plot of the pathway combined score and FDR p-value of the top 5 significant pathway scores of putative effector cells during each phase of disease.

**Supplemental Figure 8 associated with figure 6. Female mice are enriched in IFNg+ CD4 T-cells at D7 and D14**

A) Synovial flow cytometry of CD4 T-Cells at D2, D7, D14, D28 percent of CD45 (left) and live cells (right)

B) Ifng expression in synovial lymphoid cell clusters in the scRNAseq data.

**Supplemental Figure 9 associated with Figure 7. Select ISGs are sexually dimorphic in flow sorted monocytes at D2. F**low sorted monocytes (Ly6g-, Ly6c+, Cd11b+) QPCR of ISGs *Ifit, Cxcl10, Isg15 and Mx1* (A); and NfkB targets *Il1b*, *Tnf* and *Il6* (B) at D2. Each dot is one mouse, n=3-6 per group, t-test, mean and standard deviation.

**Supplemental Figure 10 associated with Figure 2. Flow gating strategy and FMO controls.**

A) Flow gating strategy to obtain target cell populations

B) Unstained and FMO controls for live cells, neutrophils, NK Cells, Cd11b+ cells, Ly6C+ cells and F4/80+ cells.

